# M2e-Derived Peptidyl and Peptide Amphiphile Micelles as Novel Influenza Vaccines

**DOI:** 10.1101/2024.06.10.598394

**Authors:** Megan C. Schulte, Agustin T. Barcellona, Xiaofei Wang, Bret D. Ulery

## Abstract

A significant problem with current influenza vaccines is their reliance on predictions of what will be the most prevalent strains for the upcoming season. Mismatches between predictions and reality in any given year can greatly reduce the overall efficacy of an immunization campaign. A universal influenza vaccine, which leverages epitopes conserved across many, if not all, strains of influenza, can reduce the need for such accurate forecasting. The ectodomain of the M2 ion channel protein is highly conserved and includes a B cell epitope in the M2_2-16_ region, making it a potentially viable candidate as a universal influenza vaccine. Unfortunately, the use of free peptide antigens as vaccines comes with several disadvantages including poor stability and weak immunogenicity *in vivo*. However, integrating peptide antigens into nanoparticles can avoid some of those drawbacks. Previous studies have shown that micellar nanoparticles can be generated from peptides by conjugating them with a lipid or lipids. Specifically, hydrophobically-driven, self-assembled peptide amphiphile micelles comprised of Palm_2_K-peptide-(KE)_4_ have been found to be immunostimulatory. Unlike other peptides previously used for this purpose, the M2_2-16_ peptide interestingly formed micelles without any peptidyl or lipid modifications. Because this unmodified peptide self-assembled on its own, it enabled the decoupling of the effect of micellization on immunogenicity from the incorporation of non-vaccine components such as the addition of a lipid moiety (Palm_2_K) and a zwitterion-like peptide block ((KE)_4_). The enclosed work shows that M2_2-16_ peptidyl micelles had some characteristic differences in shape, critical micelle concentration, and secondary structure when compared to M2_2-16_ peptide amphiphile micelles, which produced a few differences in murine antibody responses. These results suggest that peptide amphiphile micelles could be leveraged as a one-dose vaccine, while either micelle formulation induced strong immunological responses with a prime-booster immunization regimen.

## Introduction

Despite existing treatments and vaccines, influenza remains an ongoing global problem. During the 2022-2023 influenza season, there were an estimated 27 - 54 million influenza-related illnesses and 19,000 - 58,000 deaths in the United States alone.^1^ Two substantial weaknesses with our current influenza vaccine strategy employing inactivated virus are the long vaccine production times and, consequently, the need for forecasting the prevailing strains for the next influenza season. Vaccine efficacy suffers when, as a result of inaccurate predictions, the strains in the influenza vaccine do not match the dominant strains in a given influenza season.^2-5^ To address these shortcomings, there have been extensive efforts in the development of a universal influenza vaccine that can protect against a broad spectrum of influenza strains. This approach can be best achieved by harnessing conserved epitopes that are consistent across several strains of influenza. For example, the external region of the M2 ion channel protein (M2e, the first twentyfour amino acids of the protein) is one such region, with a high degree of similarity across several influenza strains including H1N1, H3N2, and H2N2.^6-8^ Excitingly, this region, more specifically M2_2-16_, has been shown to contain a B cell epitope.^9-11^

Subunit vaccines, which leverage individual antigenic components (*e*.*g*., M2_2-16_) instead of using the entire pathogen, are an attractive option to decrease vaccine production time. These antigen components can be delivered in the form of recombinant viral proteins, recombinant nucleic acids that code for these proteins, or synthetic peptides, any of which can be specifically chosen to only encode for or include the absolutely necessary immunogenic portions of the larger protein. Peptide vaccines are specifically produced via solid-phase synthesis, which is much faster than the process of growing whole virus vaccines in eggs or other vectors and do not directly compete with food or agricultural industries. Also, peptides, especially when lyophilized, have the benefit of remaining functional at room temperature for days to weeks and for years when refrigerated, with reconstitution being as simple as adding buffered solutions and mixing.^12, 13^ This stability has the potential to make peptide vaccines more accessible to underdeveloped areas with less reliable cold chain storage systems than nucleic acid or protein vaccines, which can denature in inconsistent or warmer temperatures.^12, 14^ However, peptide vaccines also have some disadvantages, including lacking inherent immunogenicity, often being unable to maintain their native conformation, and being susceptible to protease-mediated degradation in the body.^15, 16^

Many of the disadvantages of peptide vaccines can be abrogated by packaging peptides into delivery vehicles. One of the easier modes of incorporating peptides into nanoparticle-based systems is to induce peptide self-assembly, such as through micellization. Micelles are self-assembled structures that are held together by supramolecular forces including hydrophobic, ionic, and other weaker interactions. Compared to free peptides, products that form such ultrastructures are less susceptible to enzymatic degradation and have a more defined secondary structure.^17-19^ Micelles also facilitate the delivery of an increased local concentration of peptide payload to antigen presenting cells and can improve peptide-cell association.^20^ Previous work has shown that micelle size and morphology dictate their immunogenicity and these characteristics can be tuned by a number of factors including the addition of lipids and the incorporation of hydrophilic residues.^21^ Specifically, it has been shown that peptide amphiphiles (PAs) with dipalmitoyllysine on their N-terminus and a zwitterion-like charge block on their C-terminus (*i*.*e*., Palm_2_K-*antigen*-(KE)_4_) self-assemble into peptide amphiphile micelles (PAMs) that induce the highest IgG antibody titers *in vivo*.^22^ PAM immunogenicity can be further improved through the incorporation of the TLR-2 agonist adjuvant Pam_2_CSK_4_.^23^ Altogether, Palm_2_K-*antigen*-(KE)_4_ micelles co-delivered with Pam_2_CSK_4_ have the potential to be ideal carriers to be leveraged as a universal influence vaccine platform. Thus, this work sought to investigate the enhanced immunogenicity of the M2_2-16_ peptide antigen in the context of micellization and adjuvant incorporation.

## Materials and Methods

### Peptide and Peptide Amphiphile Synthesis and Purification

The TLR2 agonist Pam_2_CSK_4_ was purchased from Invivogen. The peptides M2_2-16_, M2_2-16_-(KE)_4_, and M2e (where M2_2-16_ is SLLTEVETPIRNEWG, (KE)_4_ is KEKEKEKE, and M2e is the full protein ectodomain of MSLLTEVETPIRNEWGCRCNDSSD) were synthesized using Fmoc solid-phase peptide synthesis on Sieber Amide resin using a Tetras Peptide Synthesizer. Nα-Fmoc-L-amino acids with acid-labile protecting groups on their reactive side chains were used to build the peptides studied in this research. Fmoc protecting groups were removed using 6% piperazine and 0.1 M hydroxybenzotriazole (HOBt) in dimethylformamide (DMF). Monomer couplings were done using at least 3 equivalents monomer, 6 equivalents N,N-diisopropylethylamine (DIPEA), 2.7 equivalents hexafluorophosphate benzotriazole tetramethyl uranium (HBTU), and 3 equivalents HOBt in N-methyl-2-pyrrolidone (NMP) compared to 1 molar equivalent of peptide-on-resin. After each coupling step, 5% acetic anhydride, 7% DIPEA in NMP was added to the resin to acetylate any remaining free amines. To generate PAs, a dipalmitoyllysine group (Palm_2_K) was added to the N-terminus of peptides by coupling palmitic acids to a peptide containing a non-native N-terminal deprotected Fmoc-Lys(Fmoc). For fluorophore-labeling, 5(6)- carboxyfluorescein (FAM) or 5(6)-carboxytetramethylrhodamine (TAMRA) were attached either on the N-terminus of a non-lipidated peptide or, in the case of the PA, to the deprotected ε-amine of a Lys(Dde) added to the bioactive peptide N-terminus before the inclusion of the dipalmitoyllysine group. Dde was removed with 2% hydrazine in DMF. Fluorophores were coupled to the peptide using the same protocol as amino acid addition.

Peptides were cleaved from resin using a cleavage cocktail of 2.5% each of water, phenol, triisopropylsilane, thioanisole, and ethane-1,2-dithiol in trifluoroacetic acid (TFA) for at least 2 hours, then precipitated in ether. Peptides were purified on a reverse-phase C18 column (for non-lipidated peptides) or C4 column (for lipidated peptides) in a mobile phase of water, acetonitrile, and 0.1% TFA using a gradient of increasing acetonitrile content. M2_2-16_, M2e, and Palm_2_K-M2_2-16_-(KE)_4_ eluted at approximately 35%, 35%, and 65% acetonitrile, respectively. Fluorophore-labeled FAM-M2_2-16_, Palm_2_K-K(FAM)-M2_2-16_-(KE)_4_, and TAMRA-Pam_2_CSK_4_ products eluted at approximately 40%, 60%, and 60% acetonitrile, respectively. The purified fractions were lyophilized and analyzed using mass spectrometry-controlled high-performance liquid chromatography (LC-MS) to a purity of greater than 90%. Representative LC-MS chromatograms can be found in **Figure S1**.

### Critical Micelle Concentration

Critical micelle concentration (CMC) was indirectly determined by measuring the change in fluorescence that occurs when 1,6-diphenylhexatriene (DPH) becomes encapsulated in the core of micelles. A solution of 100 mM DPH in tetrahydrofuran (THF) was prepared and then diluted 100-fold in water. The resulting solution was diluted 1,000-fold in PBS after which a serial dilution of the peptide from 100 μM to 0.92 nM was made using the 100 mM DPH in THF solution as the diluent. After a one-hour incubation in the dark, fluorescence of the serial dilutions was measured using an excitation wavelength of 350 nm and an emission wavelength of 428 nm employing a Biotek Cytation 5 spectrophotometer. The CMC for a given product was determined to be the intersection of two logarithmic regression lines on a plot of fluorescence intensity versus peptide concentration. The first line (below the CMC) was fit to at least four points where the slope is relatively flat and the second line (above the CMC) had a slope roughly ten times that of the first line, thus indicating a substantial change in fluorescence caused by DPH entrapment within the micelle core.

### Transmission Electron Microscopy

Solutions of 20 μM peptide or PA with or without 2.22 μM Pam_2_CSK_4_ were prepared in PBS from which 5 μL was pipetted onto a carbon-coated copper transmission electron microscopy (TEM) grid and incubated for 3 - 5 minutes before excess solution was wicked away using filter paper. Then, 5 μL Nano-W (2% methylamine tungstate) was pipetted on the grid and incubated for 3 - 5 minutes before being removed by wicking. Grids were imaged using a JEOL JEM-1400 Transmission Electron Microscope at 15,000x and 25,000x magnification.

### Circular Dichroism

The circular dichroism (CD) of peptide and PA solutions (250 μM) were measured from 190 nm to 250 nm with a step size of 0.1 nm on a Jasco J-1500 Circular Dichroism spectrophotometer. The data was fit to reference CD curves of poly(lysine) and poly(glutamine) to approximate α-helix, β-sheet, and random coil content of the samples. Average percentage plus/minus standard deviation of each secondary structure over three to four runs is reported.

### Förster Resonance Energy Transfer

Förster resonance energy transfer (FRET) was conducted to determine whether heterogeneous micelles were formed when products were mixed with Pam_2_CSK_4_. Solutions of 36 μM M2_2-16_ peptide (containing 0.1 μM FAM-M2_2-16_) or PA (containing 5 uM Palm_2_K-K(FAM)-M2_2-16_-(KE)_4_) with or without 4 μM TAMRA-Pam_2_CSK_4_ were prepared. The ratios of FAM-labeled peptide (or PA) to unlabeled peptide (or PA) was chosen in order to produce similar intensities between peptide and PA groups. Samples were excited at 450 nm and fluorescent emissions were measured from 475 to 700 nm using a Biotek Cytation 5 spectrophotometer.

### Murine Immunization

Each vaccination group consisted of 4 – 5 female and 3 – 4 male BALB/c mice at 10 - 11 weeks of age. Vaccines were prepared by diluting the specified amount of peptide or PA (with adjuvant as applicable) in 100 μL PBS and incubated at 4 °C overnight to allow for equilibration. Vaccines consisted of 20 nmol of peptide or PA, with or without 2.22 nmol Pam_2_CSK_4_ as outlined in **Table 1**. On day 0, mice were immunized with a subcutaneous injection in the nape of the neck. On day 14, whole blood was collected from the saphenous veins of each mouse. An equivalent second dose of vaccine was administered to the mice on day 21. On day 35, 14 days after the second vaccination, mice were sacrificed via carbon dioxide asphyxiation, following ACUC protocol guidelines. Cardiac puncture was used as a secondary means of euthanasia and for terminal blood collection.

**Table 1.** Vaccine Formulations Used for the *In Vivo* Murine Immunization Study.

| Vaccine Group | Vaccine Dosage |
| --- | --- |
| PBS |  |
| M2<sub>2-16</sub> | 20 nmol M2<sub>2-16</sub> |
| M2<sub>2-16</sub> / Pam<sub>2</sub>CSK<sub>4</sub> | 20 nmol M2<sub>2-16</sub>, 2.22 nmol Pam<sub>2</sub>CSK<sub>4</sub> |
| Palm<sub>2</sub>K-M2<sub>2-16</sub>-(KE)<sub>4</sub> | 20 nmol Palm<sub>2</sub>K-M2<sub>2-16</sub>-(KE)<sub>4</sub> |
| Palm<sub>2</sub>K-M2<sub>2-16</sub>-(KE)<sub>4</sub> / Pam<sub>2</sub>CSK<sub>4</sub> | 20 nmol Palm<sub>2</sub>K-M2<sub>2-16</sub>-(KE)<sub>4</sub>, 2.22 nmol Pam<sub>2</sub>CSK<sub>4</sub> |

where Palm<sub>2</sub>K is dipalmitoyllysine and M2<sub>2-16</sub> is the peptide sequence SLLTEVETPIRNEWG

### Serum Enzyme-Linked Immunosorbent Assay

Serum samples were generated by centrifuging whole blood at 9,400x G for 10 minutes and collecting the supernatant, after which they were frozen at -80 °C until further use. To quantify serum antibody content, enzyme-linked immunosorbent assay (ELISA) plates were coated overnight with a solution of 1.69 μM M2e peptide in carbonate buffer. The following day, plates were washed with 0.05% Tween-20 in PBS (all subsequent washes used the same wash buffer), then blocked with assay diluent (10% fetal bovine serum (FBS) in PBS) for 1 hour. Plates were sealed and incubated at 4 °C overnight. The next day, plates were washed, then incubated with a 3,000-fold dilution of secondary antibody (horseradish peroxidase (HRP)-conjugated goat anti-mouse, Invitrogen) in assay diluent for 1 hour. After washing, TMB (3,3’,5,5’ tetramethylbenzidine, Biolegend) was added to each well and allowed to incubate in the dark for 30 minutes before absorbance was measured at 650 nm using a Biotek Cytation 5 spectrophotometer to visualize HRP content. Samples were normalized across plates by subtracting the absorbance of assay-diluent only wells from the absorbance of each serum-containing well. Antibody titers were calculated as the lowest serum dilution with an absorbance at least twice that of the absorbance of the serum samples from PBS-vaccinated mice at the same serum dilution.

### Bone Marrow-Derived Dendritic Cell Generation and Assessment

Bone marrow was harvested from the femurs and tibias of 4-month-old BALB/c mice. Red blood cells were lysed using ACK lysing buffer. Bone marrow cells were cultured in complete RPMI media (10% FBS, 100 U/mL penicillin, 100 μg/mL streptomycin) with 20 ng/mL granulocyte-macrophage colony-stimulating factor (GM-CSF) to promote maturation into bone marrow-derived dendritic cells (BMDCs). After 10 days, BMDCs were plated in untreated 24-well plates and cultured with peptide or PA for 24 hours. Supernatants were collected and stored at -20 °C for later use. BMDCs were harvested and processed for flow cytometry analysis. More specifically, cells were removed from the plate, blocked with Trustain FcX anti-mouse CD16/32 antibody (Biolegend), then stained with PE/Cyaninine7 anti-mouse CD11c (Biolegend), FITC anti-mouse CD40 (Biolegend), and APC anti-mouse MHC II (Biolegend) before being fixed with 4% para-formaldehyde. Cells were analyzed on a BD LSR Fortessa flow cytometer within 48 hours after fixing. To evaluate cytokine secretion content (TNF-α, IL-1β, and IL-12 / IL-23) in the media supernatant, ELISAs were completed according to kit instructions (Biolegend).

### Statistics

One-way analysis of variance (ANOVA) and Tukey’s HSD (Honestly Significant Difference) tests were performed using GraphPad Prism software. Within a graph, groups that possess different letters have statistically significant difference in mean (p ≤ 0.05) whereas those that possess the same letter have similar means (p > 0.05).

## Results & Discussion

### M22-16 and Palm2K-M22-16-(KE)4 form small micelles at low concentrations making them ideal for vaccine applications

Purified M2_2-16_ peptide and Palm_2_K-M2_2-16_-(KE)_4_ PA were characterized for their capacity to self-assemble using a CMC assay (**Figure 1**). Surprisingly, M2_2-16_ peptide appeared to self-assemble with a CMC of 2.70 ± 1.60 μM (**Figure 1a**). These peptidyl micelles (PMs) were unexpected because previous unmodified peptide controls used in various PAM-related research (*e*.*g*., OVA_BT_ and vasoactive intestinal peptide) had not shown to possess this behavior.^21, 24^ This result was quite significant as it uniquely allowed for the decoupling of the effects that lipid presence and micellization have on peptide physical properties and immunogenicity. As expected, the Palm_2_K-M2_2-16_-(KE)_4_ PA also self-assembled, having a CMC of 0.15 ± 0.06 μM (**Figure 1b**), uncovering PAMs possess an order of magnitude lower CMC compared to analogous PMs. To probe the driving force behind PM micellization, CMCs of M2_2-16_ peptide were conducted at various salt concentrations (Milli-Q deionized, distilled water (ddH_2_O) and 10x PBS) and pHs (pH 3 and pH 13). Interestingly, altering salt concentration or pH did not appreciably affect the self-assembly of M2_2-16_ peptide with CMCs ranging from 0.56 - 5.29 μM determined under these conditions (**Figure S2**). As ion content and amino acid side chain charge greatly influences electrostatic complexation,^25, 26^ these results indicate that M2_2-16_ micellization was likely hydrophobically driven. When considering the hydropathy of the M2_2-16_ peptide and the Palm_2_K-M2_2-16_-(KE)_4_ PA (**Figure 1c**), the difference in CMCs between the PMs and PAMs becomes clearer. The hydrophobic forces involved in the self-assembly of PMs are most likely weaker than those in PAMs because the hydrophobic and hydrophilic domains are more interspersed in the peptide than the PA leading to a somewhat disordered aggregate structure (**Figure 1d**). In contrast, the PAs are strongly amphipathic with distinct regions of a very hydrophobic dipalmitoyllysine on the N-terminus and a very hydrophilic charge block on the C-terminus triggering micellization at a lower concentration in solution.^27^

**Figure 1.**
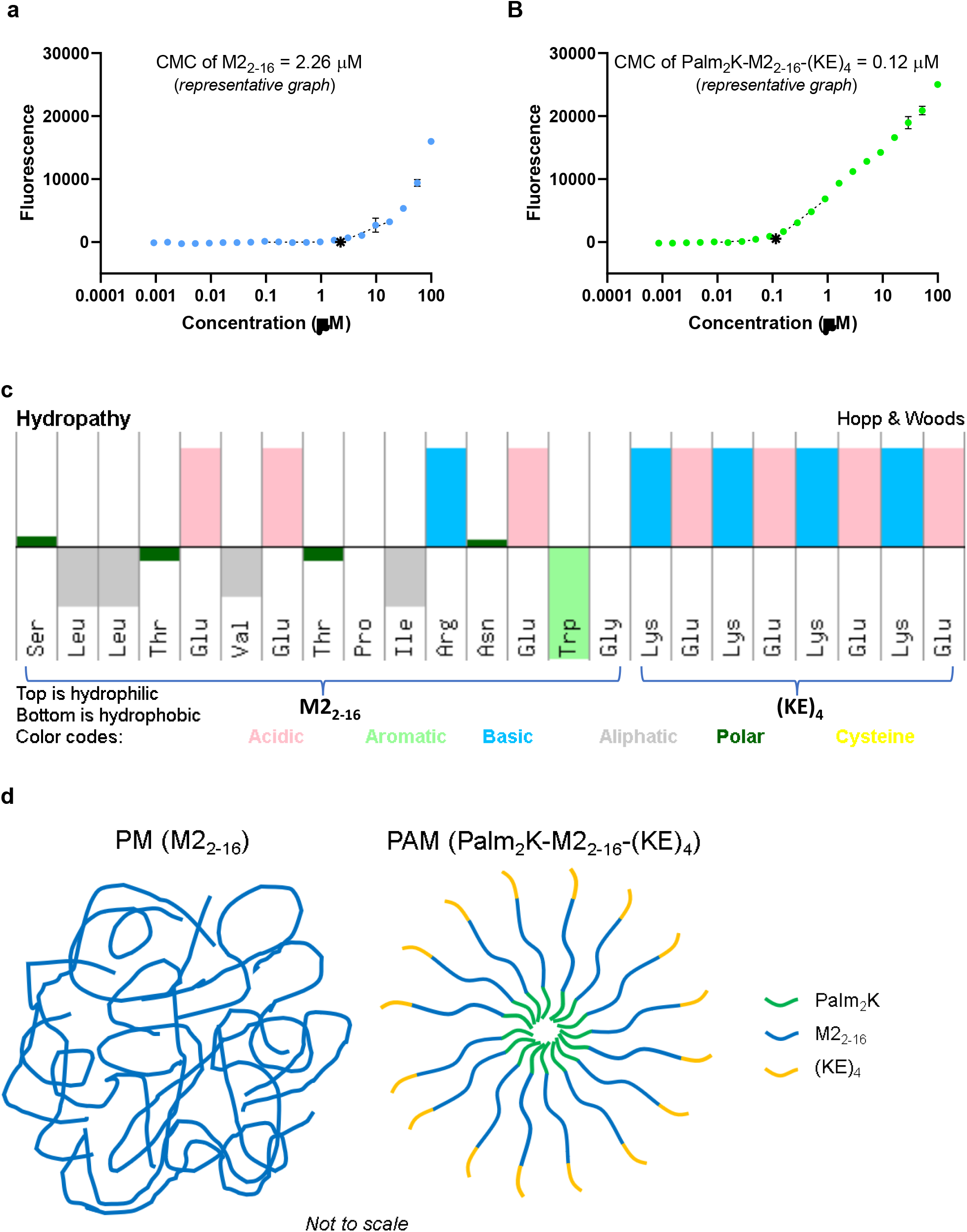
Critical micelle concentration assessment showed M2_2-16_ peptide and Palm_2_K-M2_2-16_-(KE)_4_ peptide amphiphile readily self-assemble. (a) M2_2-16_ peptide unexpectedly formed micelles at 2.26 μM (*) as shown in the representative CMC graph. (b) A representative CMC graph of Palm_2_K-M2_2-16_-(KE)_4_ PA showed micellization occurs as low as 0.12 μM (*). (c) A hydropathy chart (reprinted from Pepcalc.com) of M2_2-16_-(KE)_4_ shows that the M2_2-16_ peptide has regions of moderate hydrophobicity and hydrophilicity, whereas the (KE)_4_ region on the C-terminus contributes the most hydrophilicity to the Palm_2_K-M2_2-16_-(KE)_4_ PA.^28^ (d) With this in mind, weaker hydrophobic forces in the peptidyl micelles (left) could create more disordered micelles as compared to the distinct regions of PAs yielding more structured micelles (right).

To assess micelle morphology, both M2_2-16_ peptide and Palm_2_K-M2_2-16_-(KE)_4_ PA were visualized using TEM at a concentration above the CMC (*i*.*e*., 20 μM). Micrographs of M2_2-16_ distinctly show the formation of spherical micelles approximately 10 - 25 nm in diameter (**Figure 2a**). Interestingly, Palm_2_K-M2_2-16_-(KE)_4_ was found to produce mostly moderately short cylindrical micelles approximately 10 nm in diameter and 100 - 200 nm in length (**Figure 2b**). The modest shape difference observed between M2_2-16_ PMs and Palm_2_K-M2_2-16_-(KE)_4_ PAMs is likely a byproduct of the difference in the amphiphilicity of their constituent unimers. The more distinct hydrophobic and hydrophilic regions of PAs allow for them to pack in a more ordered, and, therefore, tighter confirmation, leading to a narrower unimer cone shape better favoring the formation of cylindrical structures over spherical ones. In contrast, the weaker definition around the hydrophobic and hydrophilic regions of the M2_2-16_ peptide likely leads to wider cones and the tendency to form spherical micelles (**Figure *1*d**).

**Figure 2.**
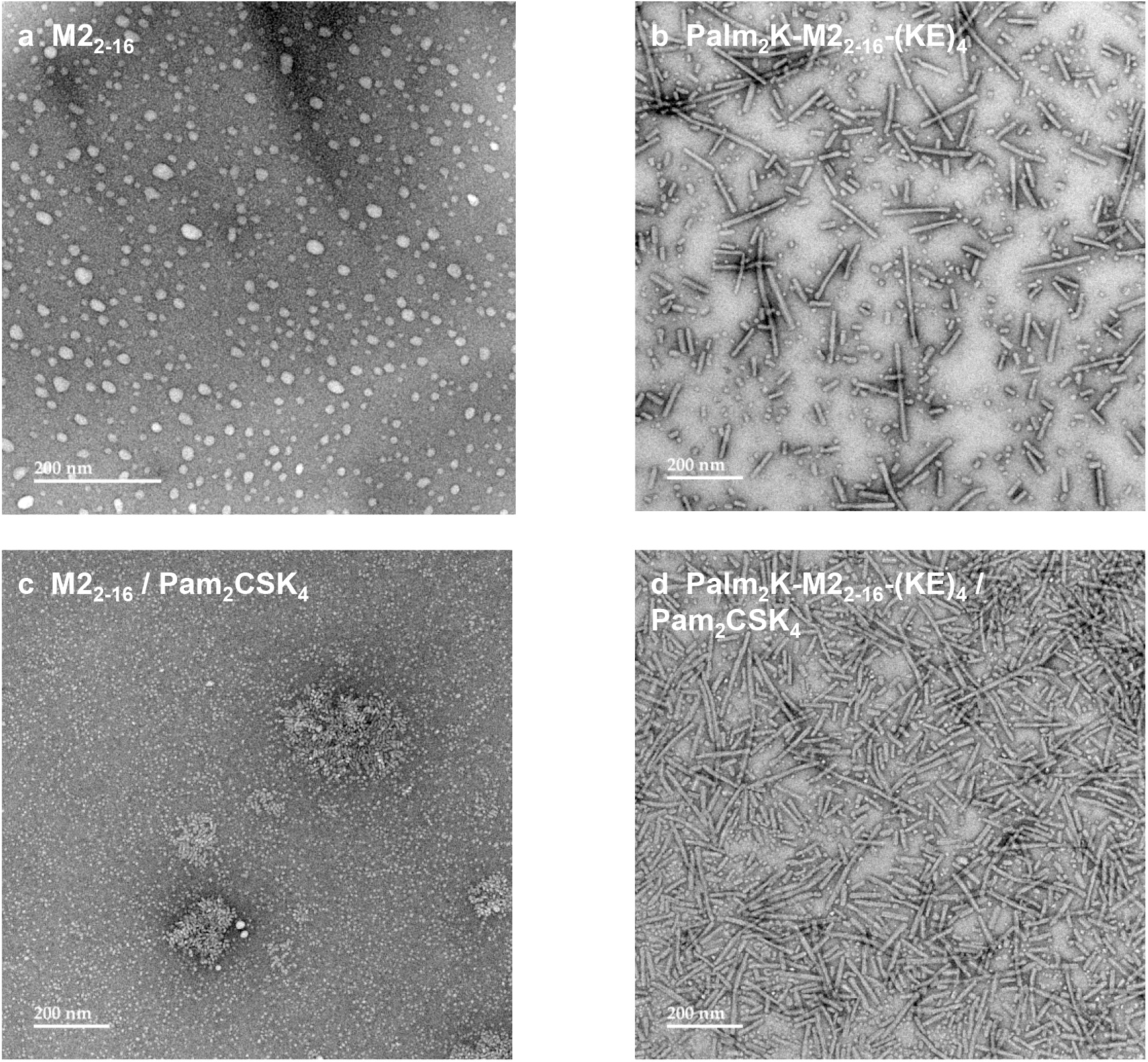
Transmission electron microscopy showed that M2_2-16_ peptide and Palm_2_K-M2_2-16_-(KE)_4_ peptide amphiphile formed small spherical and cylindrical micelles, respectively, for which the addition of Pam_2_CSK_4_ had differential effects on their morphology. (a) TEM confirmed M2_2-16_ self-assembled into small spherical micelles (∼ 10 - 25 nm in diameter). (b) Palm_2_K-M2_2-16_-(KE)_4_ formed moderately short cylindrical micelles (∼ 10 nm in diameter by 100 - 200 nm in length). (d) The addition of 10% Pam_2_CSK_4_ to 90% M2_2-16_ changed micelle morphology to smaller spheres and very short cylinders. (e) The combination of 10% Pam_2_CSK_4_ and 90% Palm_2_K-M2_2-16_-(KE)_4_ produced a mix of spherical and short cylindrical micelles, hardly impacting micelle size and shape at all.

As we were interested in studying the immunological impact of micellar adjuvant incorporation, the known TLR2 agonist and lipopeptide Pam_2_CSK_4_ was explored for its influence on PM and PAM formation. Pam_2_CSK_4_ alone was found to self-assemble into mostly small spherical micelles less than 10 nm in diameter with some occasional cylindrical micelles formed (**Figure S3**), which aligned with what has been seen in the literature.^29^ M2_2-16_ peptide or Palm_2_K-M2_2-16_-(KE)_4_ PA were then mixed with Pam_2_CSK_4_ at a 90/10 ratio and observed using TEM to assess the influence of adjuvant incorporation on micelle shape. For M2_2-16_ PMs, the presence of Pam_2_CSK_4_ eliminated the formation of micelles larger than 10 nm in diameter while preserving their spherical morphology (**Figure 2c**). In contrast, heterogeneous Palm_2_K-M2_2-16_-(KE)_4_ / Pam_2_CSK_4_ PAMs appeared to be mostly cylindrical in shape (**Figure 2d**), quite similar to Palm_2_K-M2_2-16_-(KE)_4_ PAMs alone. Even though micellization of PMs and PAMs both seemed to be hydrophobically driven, it is not entirely unsurprising that there were different responses when Pam_2_CSK_4_ was added, given the differences in amphipathicity between M2_2-16_ peptide and Palm_2_K-M2_2-16_-(KE)_4_ PA. The M2_2-16_ peptide alone does not have as strong of a hydrophobic element as the N-terminal dipalimtoyllysine of Palm_2_K-M2_2-16_-(KE)_4_, so despite the fact that M2_2-16_ PM micellization was driven by hydrophobic forces, it is possible that the morphology of the M2_2-16_ / Pam_2_CSK_4_ PMs was at least partially dictated by electrostatic forces. Specifically, the change in micelle size from M2_2-16_ to M2_2-16_ / Pam_2_CSK_4_ could be explained by the repulsion of negatively charged residues in M2_2-16_ peptide being disrupted and replaced by tight complexation with the positively charged lysines in Pam_2_CSK_4_, thus resulting in the formation of smaller micelles.^30^ In the case of the PAMs, however, Palm_2_K-M2_2-16_-(KE)_4_ and Pam_2_CSK_4_ are structurally similar as both are lipopeptides with two 16-carbon tails and charged residues on their C-terminus. Therefore, it is unsurprising that there were no substantial morphological changes to Palm_2_K-M2_2-16_-(KE)_4_ PAMs with the addition of Pam_2_CSK_4_.

To confirm that Pam_2_CSK_4_ integrated into the PMs and PAMs, FAM-labeled peptide or PA was mixed with TAMRA-labeled Pam_2_CSK_4_. The fluorophores FAM and TAMRA are a FRET pair, where the energy emitted by FAM excites TAMRA, allowing for TAMRA emission to be observed when in close physical proximity to FAM. Excitingly, FRET was clearly observed in the PAMs, as a decrease in fluorescence intensity at 525 nm (*i*.*e*., the FAM emission wavelength) and the appearance of a fluorescence local maximum at 580 nm (*i*.*e*., the TAMRA emission wavelength) were seen when TAMRA-Pam_2_CSK_4_ was present, confirming the presence of heterogeneous micelles (**Figure 3**). While the M2_2-16_ PMs did not exhibit a fluorescence maximum at 580 nm, there was a decrease in fluorescence intensity at 525 nm when in the presence of TAMRA-Pam_2_CSK_4_, which suggested that there was FRET and, consequently, that M2_2-16_ and Pam_2_CSK_4_ formed heterogenous micelles. To provide further evidence, the fluorescence of increasing ratios of TAMRA-Pam_2_CSK_4_ to M2_2-16_ peptide was measured. With higher concentrations of TAMRA-Pam_2_CSK_4_ a local maximum at 580 nm was observed, validating the existence of FRET in the PMs (**Figure S4**).

**Figure 3.**
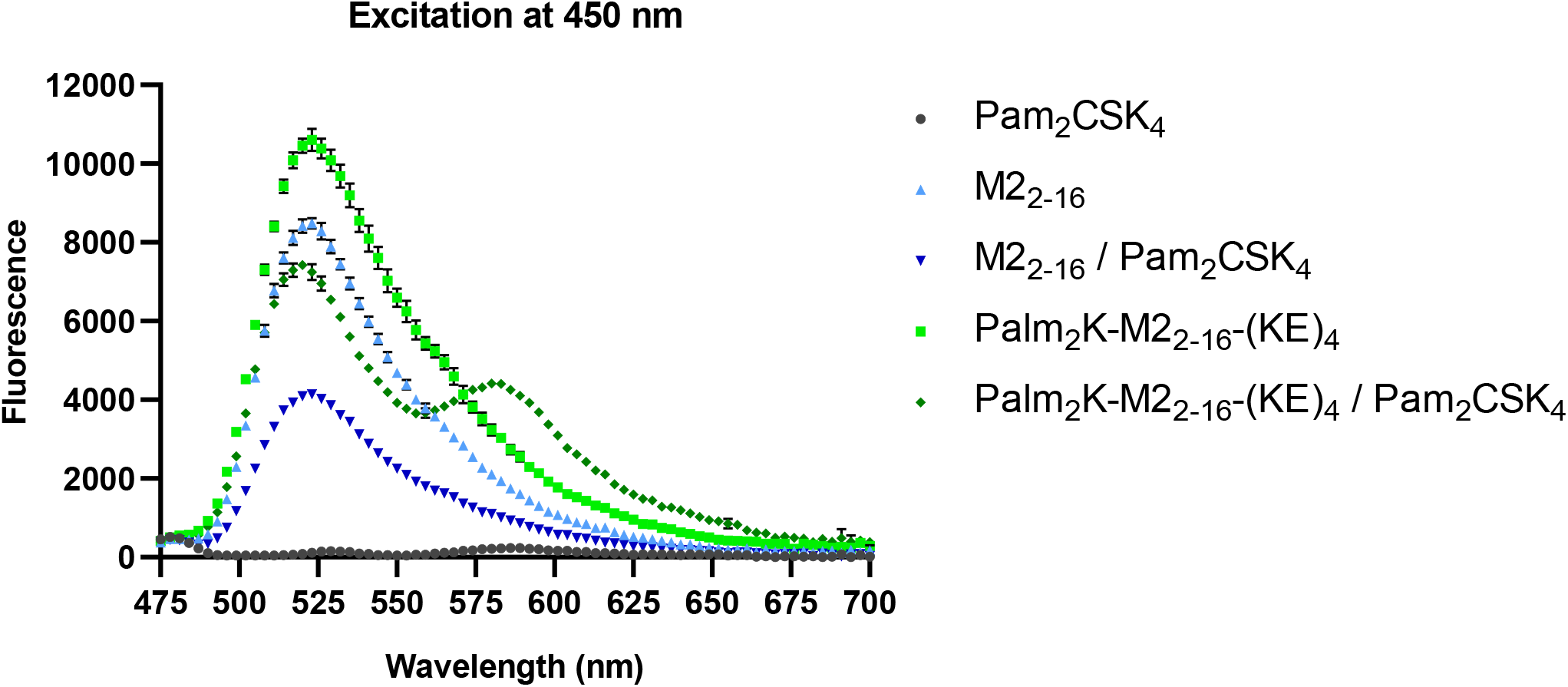
Changes to the fluorescence spectra at 525 nm of the peptides and peptide amphiphiles after adding Pam_2_CSK_4_ confirmed that Pam_2_CSK_4_ was integrated into both peptidyl micelles and peptide amphiphile micelles.

Given the morphological differences between M2_2-16_ PMs and Palm_2_K-M2_2-16_-(KE)_4_ PAMs, circular dichroism (CD) analysis was carried out to evaluate whether those differences influenced peptide secondary structure (**Figure 4**). When fit to known spectra, PMs were found to be mostly random coil (61.3%) in nature, as shown by a minimum at 198 nm, but they also had considerable β-sheet character (38.7%). PAMs, on the other hand, were determined to be almost entirely β-sheet (i.e., 97.5%), as shown by a minimum at 219 nm and a maximum at 205 nm. The structure of the M2 protein ectodomain in the context of the influenza virus has not been well characterized, except when untethered from the rest of the protein and bound to antibodies or other entities. Results in the literature and using predictive software (PEP-FOLD and I-TASSER) have been inconsistent, showing the presence of some α-helix or β-sheet or only random coil. ^31-34^ Although there was not a clear expectation of what a desired secondary structure composition would be for the micelles, Qiao, *et al*., from an investigation of 350 antigen-antibody complexes, found that polar regions with more constrained secondary structure (*e*.*g*. helices or sheets) on the surface of a protein were more favorable for antibody recognition.^35^

**Figure 4.**
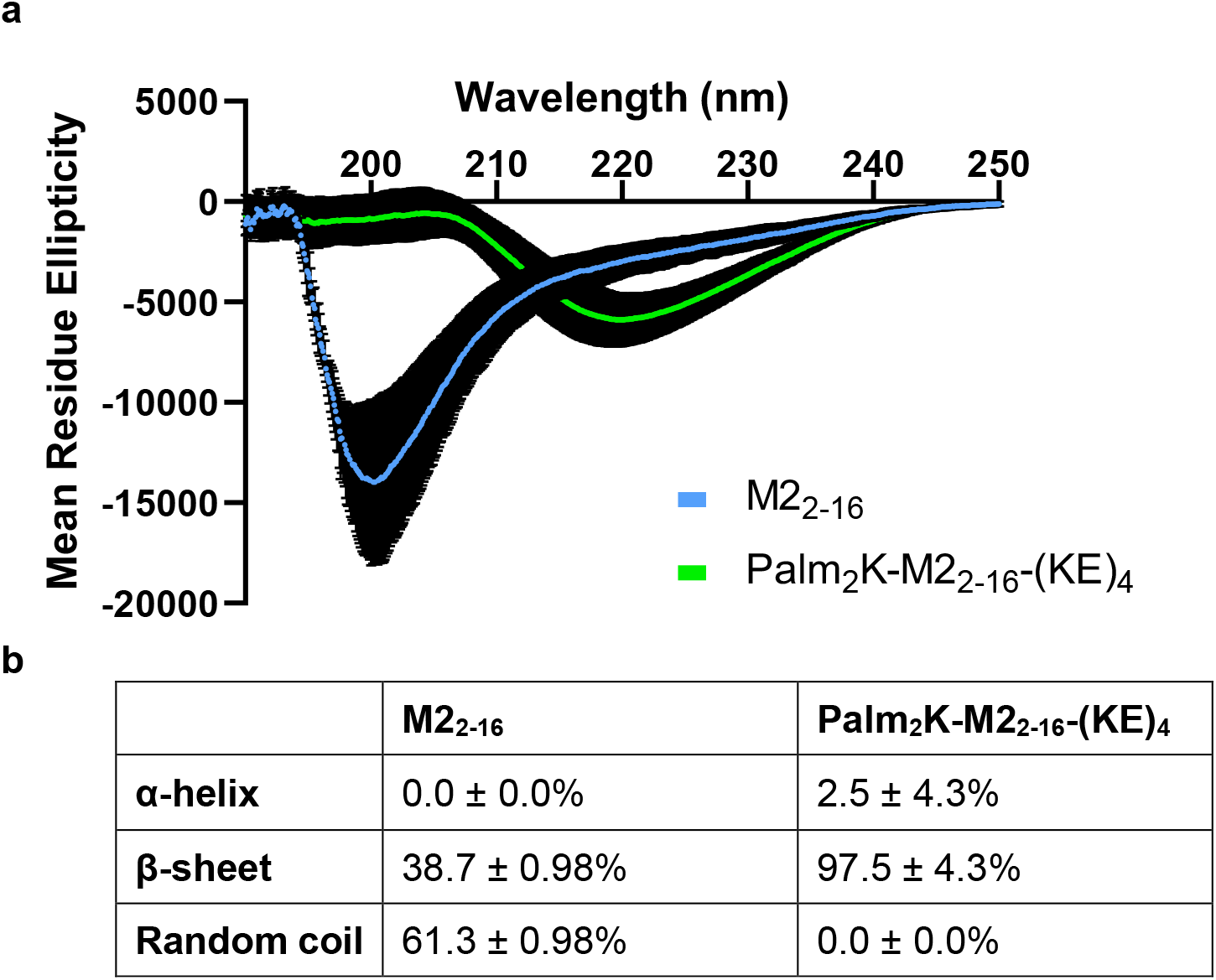
Peptidyl micelles and peptide amphiphile micelles had distinctly different secondary structures. (a) Circular dichroism (CD) spectra of M2_2-16_ PMs and Palm_2_K-M2_2-16_- (KE)_4_ PAMs showed minima at 200 nm and 219 nm, respectively, and Palm_2_K-M2_2-16_-(KE)_4_ PAMs has a maxima at 204 nm. (b) Based on reference CD data for poly(lysine) and poly(glutamine), the estimated secondary structure of M2_2-16_ PMs was mostly random coil whereas Palm_2_K-M2_2- 16_-(KE)_4_ PAMs was almost entirely β-sheet.

### Peptide amphiphile micelles elicited strong antibody titers after the primary immunization, but both micelle types elicited strong IgG titers after the booster

To evaluate the immune response against both micelle types, mice were subcutaneously vaccinated with both formulations (with and without the adjuvant Pam__2__CSK__4__). Blood was collected at days 14 and 35 (14 days after the first and second vaccinations, respectively). ELISAs were conducted on the serum from the blood collections to evaluate antibody production against the vaccines. Interestingly, at day 14, there were no IgM responders in either PM groups (with or without adjuvant), but both PAM groups had similar statistically significantly higher titers over the PBS vaccine baseline, independent of the presence of Pam_2_CSK_4_ (**Figure 5a**). By day 35, although there was no longer statistical differences between any groups, there were still several more responders in the PAM vaccine groups (10 total) compared to the PM vaccine groups (3 total), although PAM IgM titers did seem to have decreased from day 14 (**Figure 5b**). Day 14 IgG titers showed a few responders in the mice vaccinated with M2_2-16_, M2_2-16_ / Pam_2_CSK_4_, and Palm_2_K-M2_2-16_-(KE)_4_, but when combined with non-responders, these results were found to be statistically insignificantly above background (**Figure 5c**). Excitingly, however, the adjuvant-loaded PAM formulation (*i*.*e*., Palm_2_K-M2_2-16_-(KE)_4_ / Pam_2_CSK_4_) induced statistically appreciable IgG content at day 14, suggesting its potential as a single-dose vaccine. This would be especially relevant for vaccines against seasonal or rapidly emerging pathogens like influenza and COVID-19, respectively. The day 35 IgG data showed that both adjuvant-free micelles (*i*.*e*., M2_2-16_ and Palm_2_K-M2_2-16_-(KE)_4_) induced an antibody response though interestingly PMs induced slightly higher, though statistically insignificant, titers above PAMs (**Figure 5d**). The incorporation of adjuvant in either PMs or PAMs yielded vaccines that induced substantially elevated IgG titers in which differences between the micelle-type were minimized.

**Figure 5.**
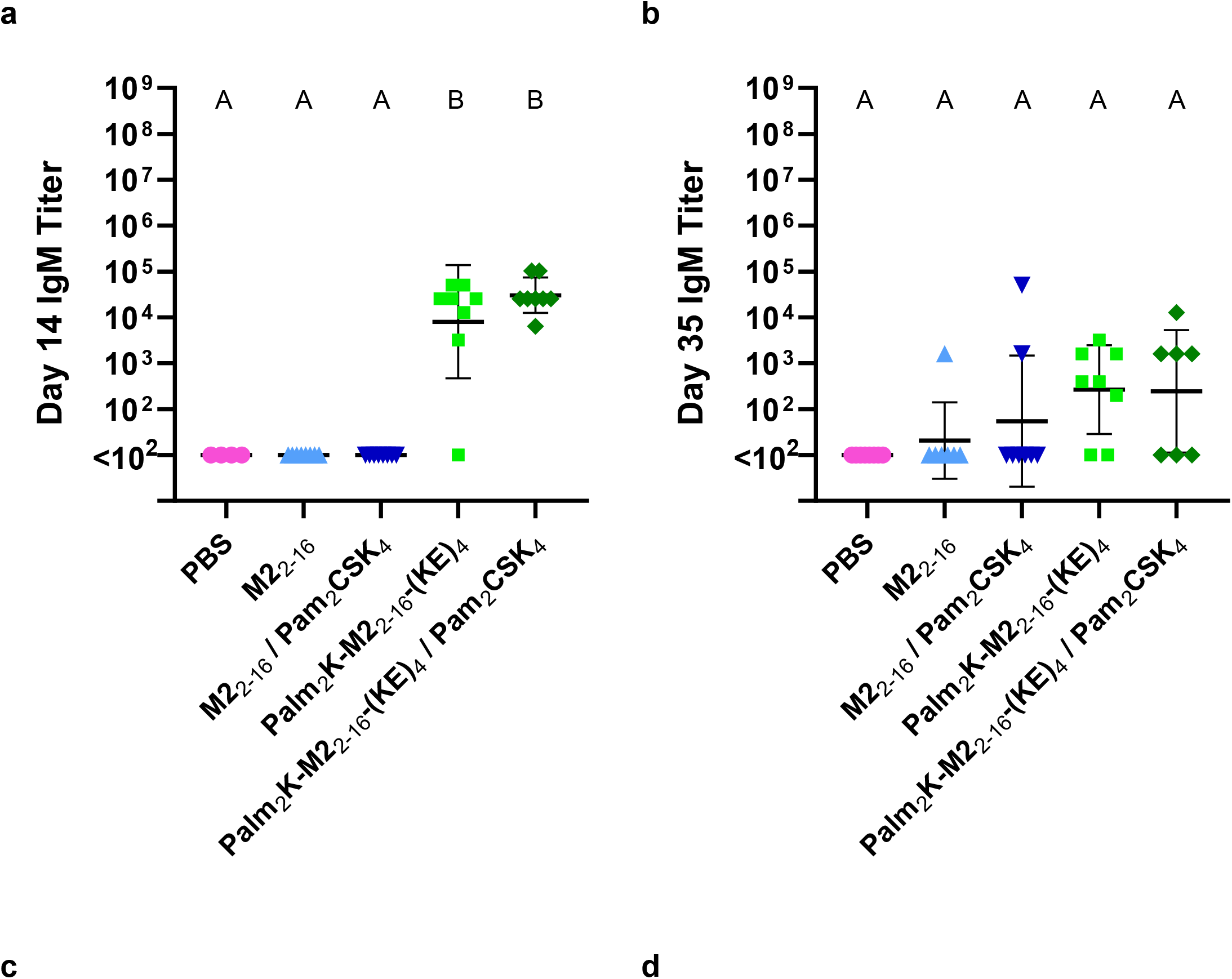
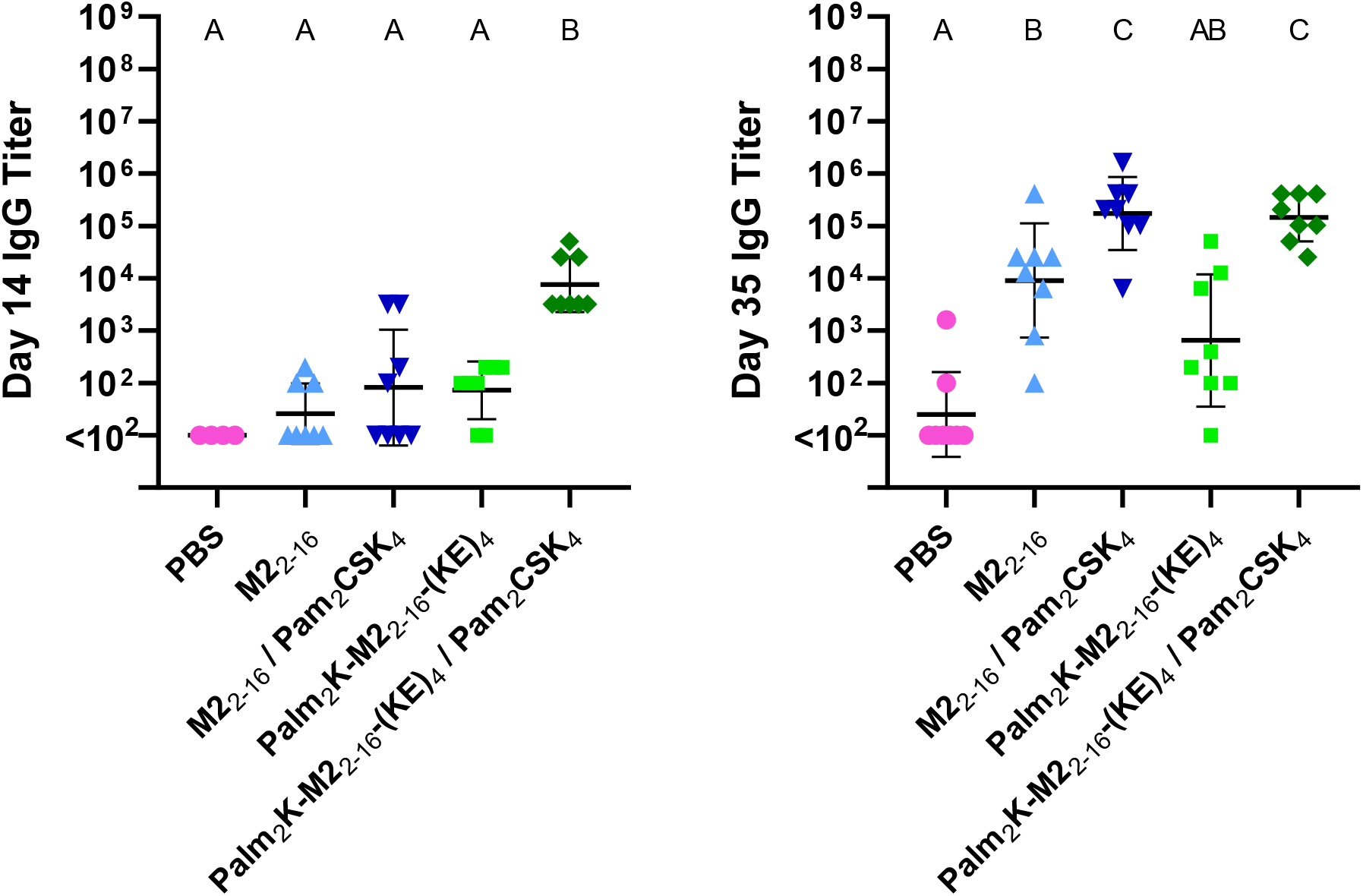
Serum titers from peptide amphiphile micelle-vaccinated mice were the strongest at day 14, while later responses were primarily driven by co-delivery with Pam_2_CSK_4_. (a) PAMs induced a strong IgM response at day 14, regardless of adjuvant. (b) No significant difference was found in IgM titers between any vaccine groups in day 35 serum, although PAMs were slightly higher than background though less than what was observed at day 14. (c) Only Palm_2_K-M2_2-16_-(KE)_4_ / Pam_2_CSK_4_ induced IgG titers at day 14 above baseline. (d) All micelle groups elicited IgG titers above baseline in day 35 serum, though Palm_2_K-M2_2-16_-(KE)_4_ was unable to do so with statistical significance. Pam_2_CSK_4_-containing groups elicited statistically significantly elevated titers over groups without Pam_2_CSK_4_. Within a graph, groups that possess different letters have statistically significant difference in mean (p ≤ 0.05) whereas those that possess the same letter have similar means (p > 0.05).

The day 35 IgG titers seen here differed from similar studies previously conducted with different antigens.^23,36^ In those studies, target antibody titers in mice vaccinated with non-micellized peptide alone were low. Mice vaccinated with peptide and adjuvant or PAM without adjuvant had intermediate titers and vaccinations of PAM with adjuvant produced the highest titers. There are several hypotheses for why the results of this study differed from previous studies, namely why the PAMs did not produce higher titers than the peptides. Firstly, micellization of the unmodified peptide antigen was not seen in previous cases, so it is possible that the micellization of the M2_2-16_ peptide elicited higher antibodies compared to a non-micellized peptide control – this would make the PAM titers seem lower when, in reality, the “control” was just comparatively higher. Another possibility is that the PAMs were still generating an overall higher titer, but that some of the antibodies were not being captured in the ELISA (*e*.*g*. because these antibodies were not specific to the M2_1-24_ antigen). The relatively higher IgM titers of the PAMs compared to the peptidyl micelles could be evidence of this effect. Investigating these hypotheses was outside the scope of this work, but should be explored in the future. Regardless, it is clear that PAMs produced a superior initial antibody response to the first dose of the vaccine compared to PMs, making PAMs a good candidate for single-dose vaccines. Even after the second dose, the antibody responses between PMs and PAMs were comparable, suggesting a potential for uses in multi-dose vaccines, especially in cases where delivering boosters might not always be possible, such as in rural areas without easy access to healthcare.

To further investigate the antibody response, ELISAs were conducted on the serum collected from day 35 to measure the relative amounts of the IgG subclasses IgG1, IgG2a, and IgG3 present (**Figure 6**). IgG1 titers (**Figure 6a**) largely aligned with total IgG titers (**Figure 5d**), in that titers were highest in Pam_2_CSK_4_-containing vaccine groups and that the PMs had a slightly higher (although statistically insignificant) titer than the PAMs. The PAMs without any adjuvant did not produce a titer statistically above baseline. For IgG2a titers, only the PMs with adjuvant and PAMs with adjuvant elicited titers statistically above baseline (and the majority of serum samples in the groups without adjuvant were non-responders) (**Figure 6b**). IgG3 titers largely aligned with IgG2a titers, except that there were more positive responders in all groups (**Figure 6c**).

**Figure 6.**
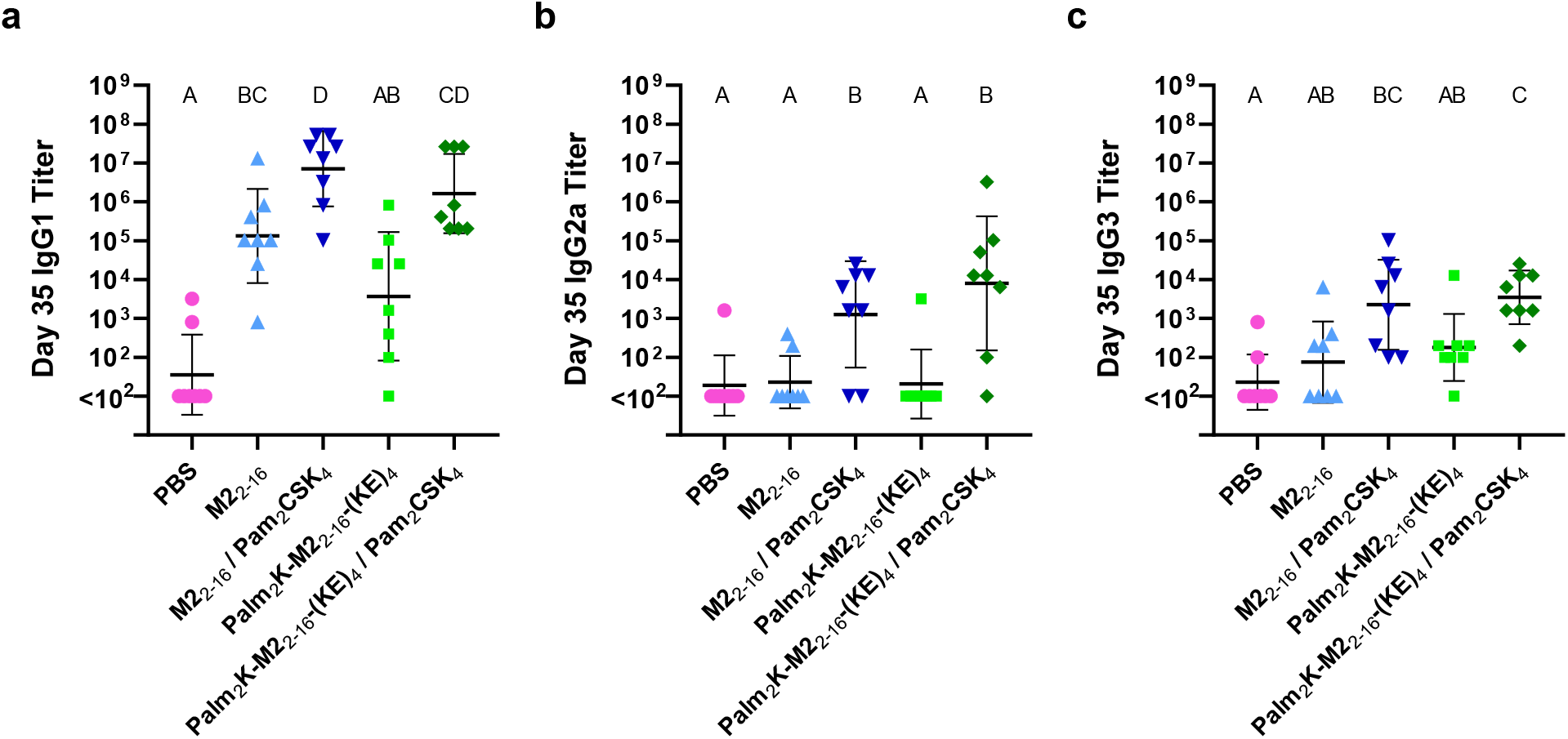
Immunoglobulin subtype assessment showed an IgG1-skewed antibody response for peptidyl and peptide amphiphile micelles. (a) IgG1 titers were highest in Pam_2_CSK_4_-containing vaccine groups, with the M2_2-16_ peptidyl micelles having slightly (not significant) higher titers than Palm_2_K-M2_2-16_-(KE)_4_ peptide amphiphile micelles. (b) IgG2a titers were lower in number than IgG1, with only adjuvant-containing groups above baseline. Although not statistically significant, Palm_2_K-M2_2-16_-(KE)_4_ / Pam_2_CSK_4_ had a slightly higher titer than M2_2-16_ / Pam_2_CSK_4_. (c) IgG3 titers matched the IgG1 titers trend though with much lower antibody titers. Micelles without adjuvant were above baseline, but not statistically significantly. Micelles with adjuvant produced comparable titers above baseline, but only Palm_2_K-M2_2-16_-(KE)_4_ / Pam_2_CSK_4_ was statistically significantly above micelles without adjuvant. Within a graph, groups that possess different letters have statistically significant difference in mean (p ≤ 0.05) whereas those that possess the same letter have similar means (p > 0.05).

While the direct ratiometric comparison of titers across subclasses is not perfect because of the use of different secondary antibodies that might have different binding characteristics, it is evident, especially in the formulations without adjuvant, that IgG1 was the primary subclass generated in response to all vaccine formulations, with lower responses in IgG2a and IgG3. This aligned with several studies that used PAMs containing different antigens – IgG1 was the dominant subclass and IgG2a titers were low.^23, 36, 37^ An ideal influenza vaccine response would mimic the natural host response to better prepare the immune system for exposure to the influenza virus. In mice, IgG2a is the predominant antibody generated in the immune response against influenza infection, followed by IgG1, then IgG3.^38^ While the IgG subtype ratio was not IgG2a-dominant in this study, the administration route of the vaccine could have played a role in the outcome. Wareing, *et al*. compared the generation of IgA and IgG2a antibodies for an influenza vaccine via different routes of administration: subcutaneous, intramuscular, and intranasal.^39^ They found that intranasal delivery produced the highest IgA antibodies in the lungs and the highest serum IgG2a. To our knowledge, PAM vaccines have not been administered intranasally to date, but this could be a worthwhile approach to produce a more desired antibody profile in a future study.

### Bone marrow-derived dendritic cells were activated by adjuvant-supplemented peptidyl micelles and peptide amphiphile micelles

To test the innate immunogenicity of the micelle formulations, bone marrow-derived dendritic cells (BMDCs) were cultured and treated with 1.8 μM antigen-containing peptide (PMs or PAMs) with or without 0.2 μM Pam_2_CSK_4_ for 24 hours. BMDCs were evaluated using flow cytometry and identified by CD11c expression (**Figure S5**). BMDC activation was identified by elevated CD40 and MHC-II expression. The percent of CD11c^+^ cells expressing CD40 was significantly increased among cells exposed to all treatments containing Pam_2_CSK_4_ (*i*.*e*. Pam_2_CSK_4_ alone and with either micelle type) without measurable differences between those groups (**Figure 7a**). There was less of an effect (not statistically significant) for MHC-II expression, although the mean cell percentages for CD11c^+^MHC-II^+^ cells in the Pam_2_CSK_4_-treated groups were higher than in groups not treated with Pam_2_CSK_4_. The adjuvant effect was still present when comparing cell numbers between peptide treatments in the CD11c^+^ population expressing both activation markers (CD40 and MHC-II), although this was likely a carry-over effect from the significant differences between groups in the CD40^+^CD11c^+^ population. When looking at the median fluorescence intensity (MFI) of the FITC-CD40 signal even among CD40^+^CD11c^+^ cells, there was again a distinct and similar signal enhancement within Pam_2_CSK_4_-treated groups compared to groups treated with PMs or PAMs alone (**Figure 7b**). This enhancement was also seen in the MFI of the APC-MHC-II signal of MHC-II^+^CD11c^+^ cells (despite there not being any significant effect on MHC-II expression based on cell numbers). When looking at the MFI of FITC-CD40 and APC-MHC-II among CD40^+^MHC-II^+^-CD11c^+^ cells, there was only a difference between treatment groups within the FITC-CD40 channel, again following the pattern of Pam_2_CSK_4_-treated *vs*. not Pam_2_CSK_4_-treated (**Figure 7c**). Altogether, it is evident that BMDC activation was primarily driven by Pam_2_CSK_4_ (and, consequently, TLR-2 stimulation) and that the overall enhancement of CD40 expression was more pronounced than MHC-II.

**Figure 7.**
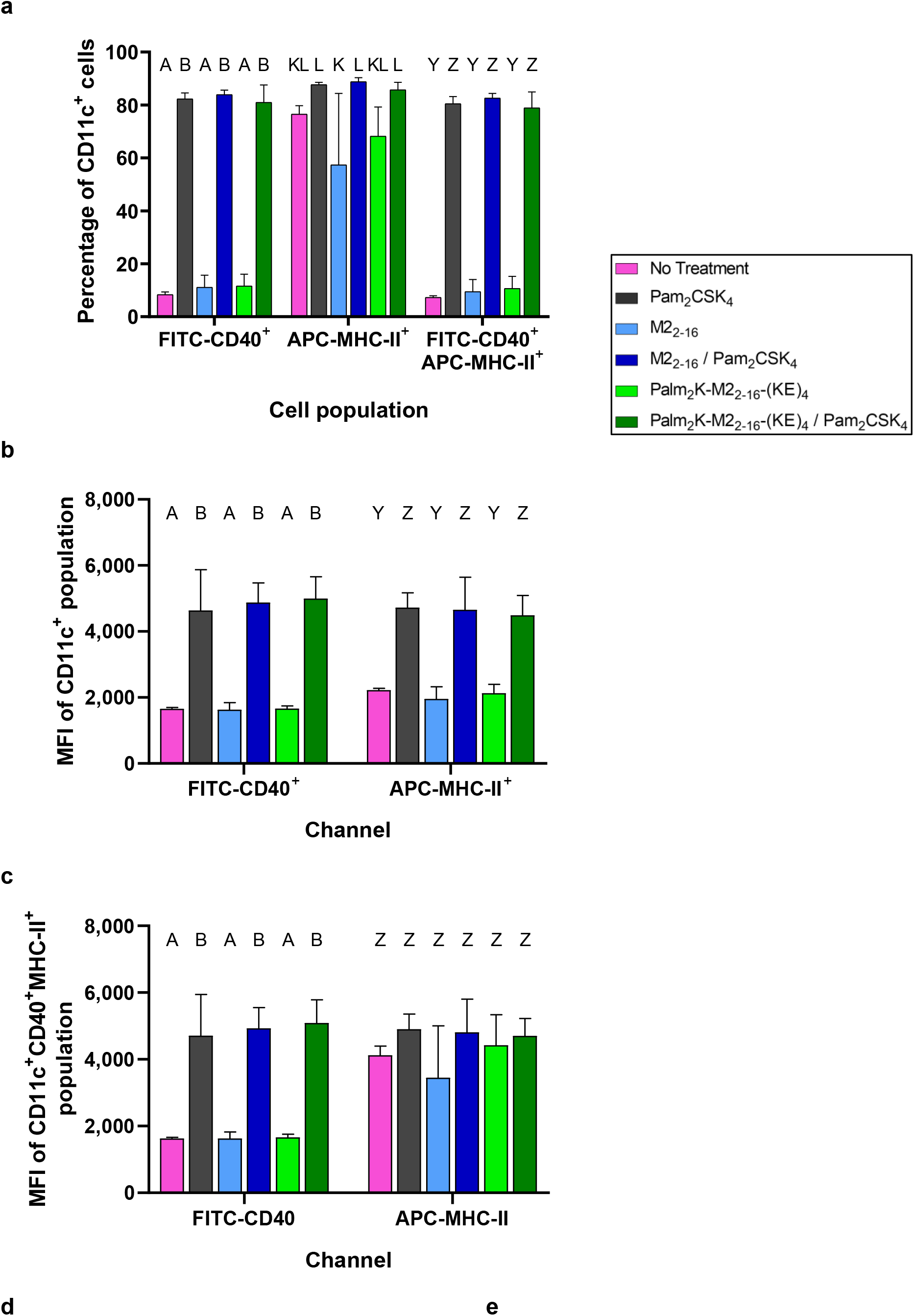
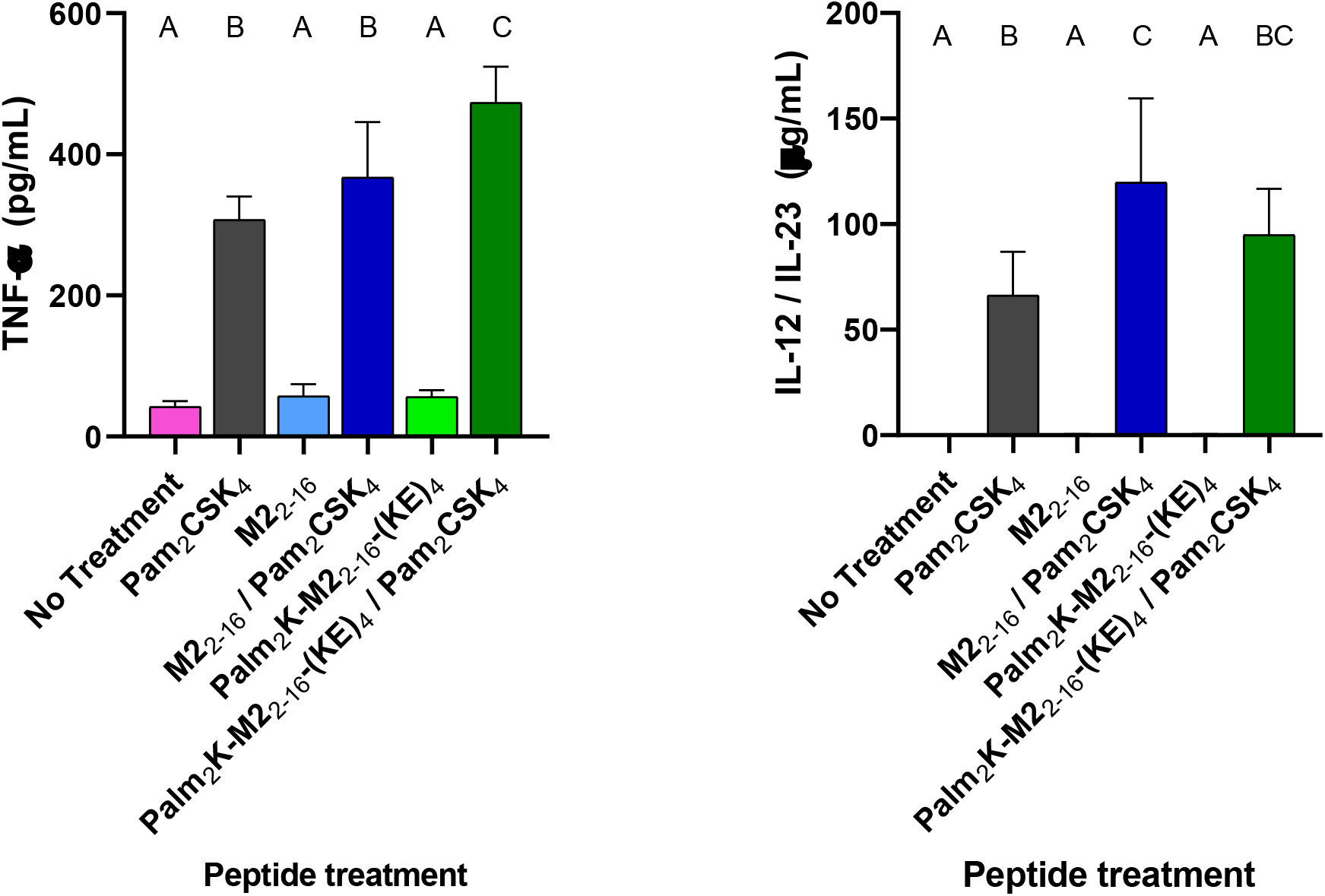
Bone marrow-derived dendritic cell activation is primarily affected by the presence of Pam_
2_CSK_4._ (a) The percentage of CD11c^+^ cells expressing CD40 (but not MHC-II) increased significantly when treated with Pam_2_CSK_4_-containing formulations. (b) Treatments containing Pam_2_CSK_4_ strengthened the MFIs of the FITC and APC channels for CD11c^+^CD40^+^ and CD11c^+^MHC-II^+^ cell populations, respectively. (c) The MFI of CD11c^+^CD40^+^MHC-II^+^ was elevated in the FITC channel, but not the APC channel for Pam_2_CSK_4_-containing treatments. Pam_2_CSK_4_ elicited higher production of proinflammatory cytokines (d) TNF-α and (e) IL-12 / IL-23, as determined by ELISAs. Groups that possess different letters have statistically significant difference in mean (p ≤ 0.05) whereas those that possess the same letter have similar means (p > 0.05). Statistical comparisons were only made between different peptide treatments within the same channel or gating scheme.

Sandwich ELISAs were conducted on supernatant media collected from treated BMDCs to evaluate their secretion of the pro-inflammatory cytokines IL-1β, TNF-α, and IL-12 / IL-23 (by quantifying the subunit p40) . No group from the IL-1β ELISA was consistently above the detection limit of 62.5 pg/ml (*data not shown*). IL-1β has previously been shown to be upregulated with TLR-2 activation by di-or tri-acylated lipopeptides, so it is possible that the dose of Pam_2_CSK_4_ was too low to observe any changes in IL-1β secretion or that co-stimulation of another receptor was needed.^40, 41^ In contrast, TNF-α and IL-12 / IL-23 secretion in BMDCs (**Figure 7d & 7e**) were most largely impacted by the presence of Pam_2_CSK_4_, as was seen in the flow cytometry data. Pam_2_CSK_4_ is a known TLR-2 agonist, which triggers TNF-α production via the TLR-2-MyD88-NF-κB pathway.^42, 43^ IL-12 has similarly been shown to be highly expressed with TLR-2 activation in BMDCs.^44-46^ Given the role of dendritic cells in antigen presentation and lymphocyte activation, it is likely that there is a direct relationship between the BMDC activation data and antibody titer data.

## Conclusion

Compared to what has been observed in other investigations utilizing PAMs, the M2_2-16_ antigen micellized, which enabled the unique opportunity to decouple the effects of micellization and antigen modification on the immune response of a micellized peptide vaccine. As has been previously demonstrated in literature using other peptides, the addition of an N-terminal dipalmitoyllysine and a C-terminal charge block to the M2_2-16_ peptide produced short cylindrical micelles with increased β-sheet character and lower CMCs compared to the unmodified M2_2-16_ peptidyl micelles. The changes in physical characteristics seemed to induce slight changes in the immune response. Although the overall IgG response was comparable between PMs and PAMs, the PAM formulation interestingly elicited significantly higher IgM and IgG titers in the initial antibody response at day 14 and even slightly (although statistically insignificant) higher IgM titers than the PMs at day 35. While the PMs seemed to produce a slightly improved immune response in the context of the prime-booster vaccine regimen, it appears that PAMs could be better candidates for single-dose vaccines.

## Supporting information

Full Supplemental Information

## Author Contributions

**Megan C. Schulte** – writing, editing, experimental design, experimental work (characterization, *in vitro* work, *in vivo* work)

**Agustin Barcellona** – experimental work (*in vivo* work)

**Xiaofei Wang** – experimental work (*in vivo* work)

**Bret D. Ulery** – writing, editing, conceptualization, funding acquisition

## Acknowledgments

The authors would like to thank the Molecular Interactions Core at the University of Missouri for use of the Tetras Peptide Synthesizer, LC-MS, and J-1500 circular dichroism spectrophotometer, especially Fabio Gallazzi for his assistance with peptide synthesis. The authors would also like to thank the Electron Microscopy Core at the University of Missouri for use of the JEOL JEM-1400 Transmission Electron Microscope.

