## Supplementary material for "M2e-Derived Peptidyl and Peptide Amphiphile Micelles as Novel Influenza Vaccines": Full Supplemental Information

### Supplementary Information

**a**

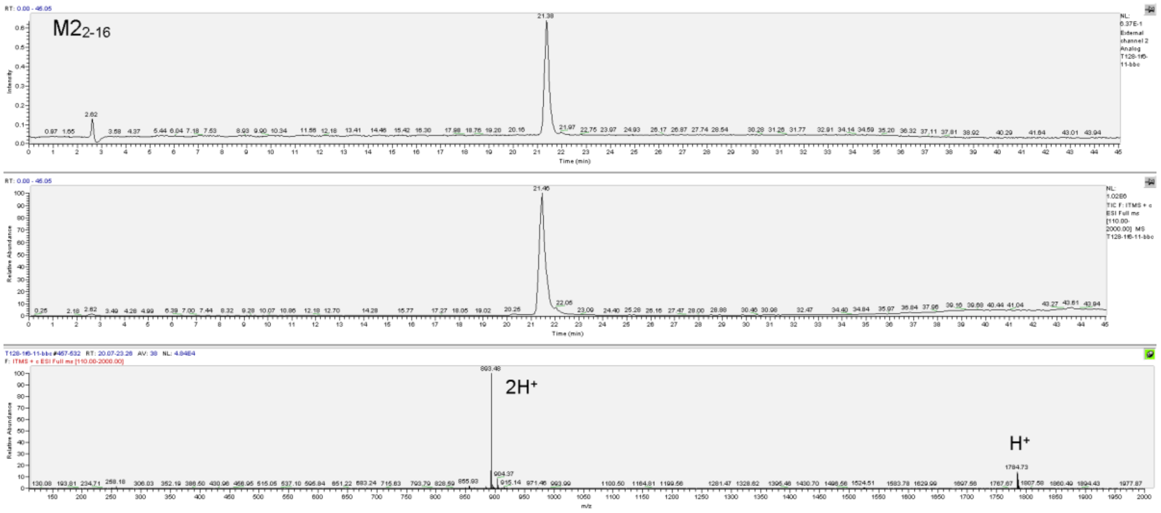


**b**

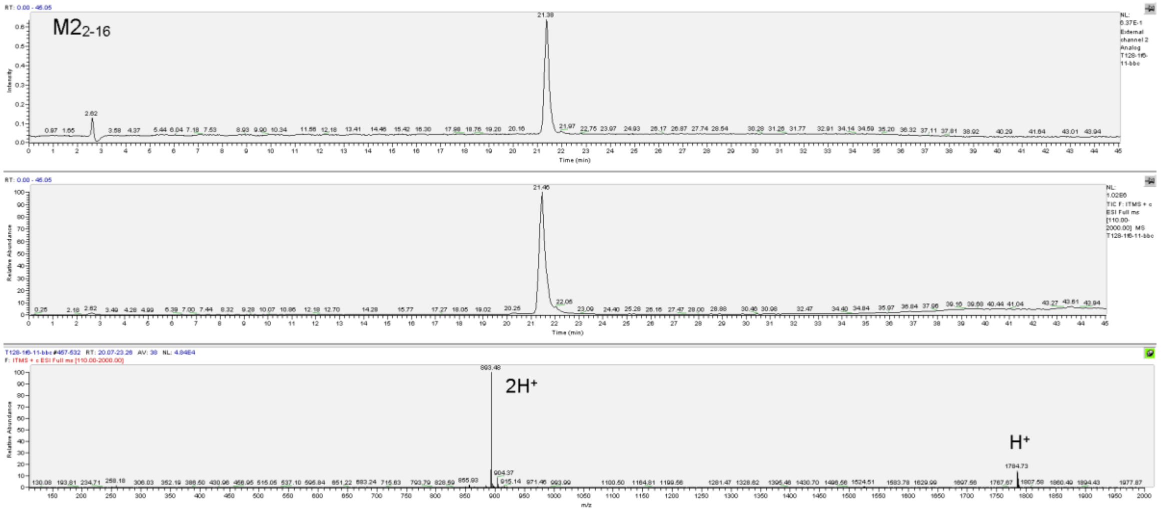

**c**

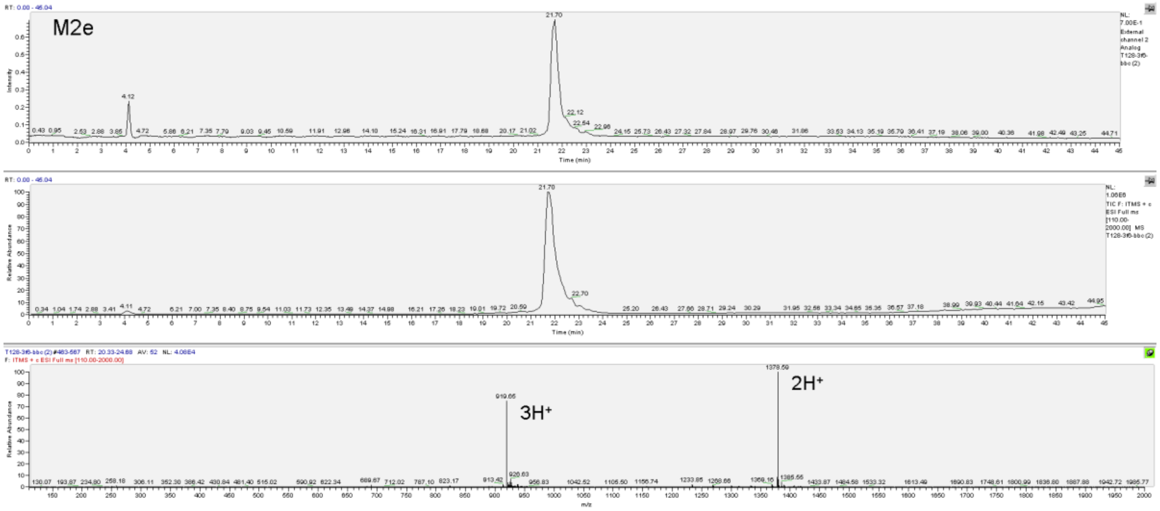
M2e
Time (min)
3H+
2H+
m/z

**d**

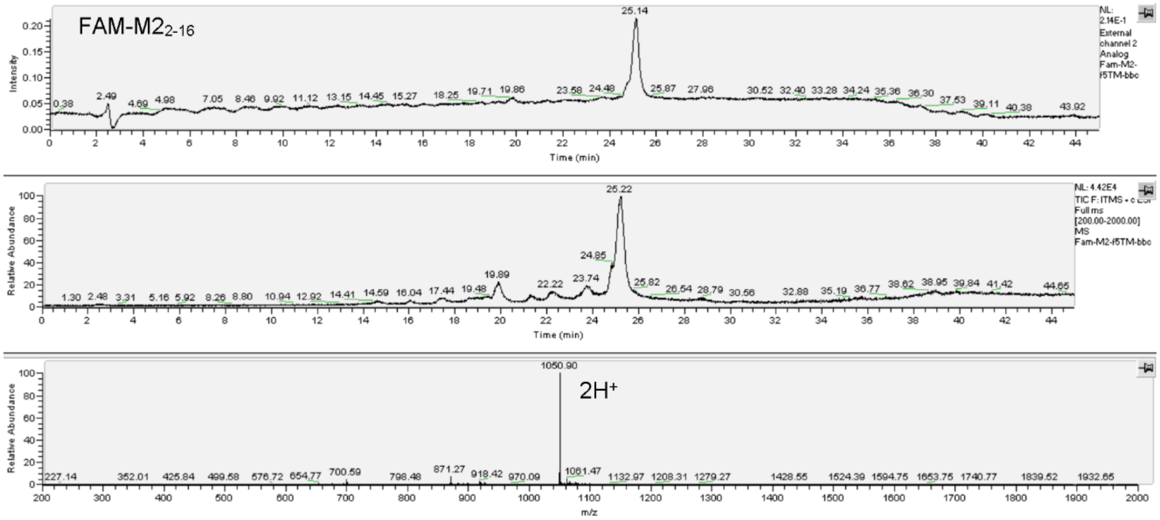
FAM-M22-16
Intensity
Time (min)
Relative Abundance
2H+
m/z

e

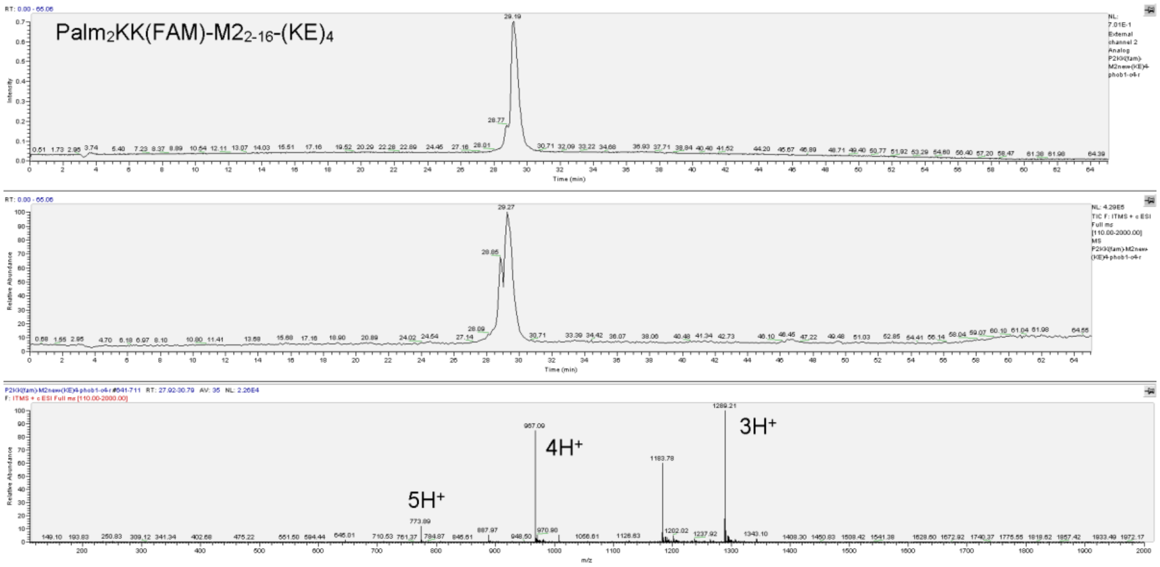


f

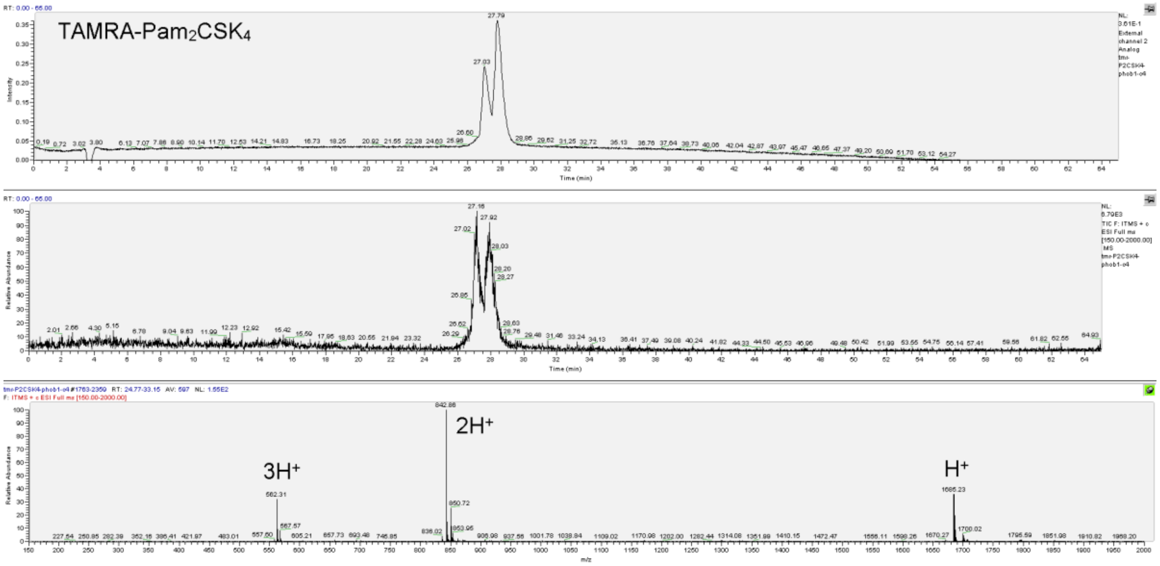


**Figure S1. Peptides were purified to greater than 90% purity using LC-MS.** LC-MS analyses are shown for purified (a) $M2_{2\text{-}16}$, (b) $Palm_2K\text{-}M2_{2\text{-}16}\text{-}(KE)_4$, (c) M2e, (d) $FAM\text{-}M2_{2\text{-}16}$, (e) $Palm_2K\text{-}K(FAM)\text{-}M2_{2\text{-}16}\text{-}(KE)_4$, and (f) $TAMRA\text{-}Pam_2CSK_4$. The top, middle, and bottom figures of each panel consist of a UV chromatograph, a total ion count, and a mass spectra, respectively, For the mass spectra, splitting due to single through quadruple charge states are denoted where applicable.

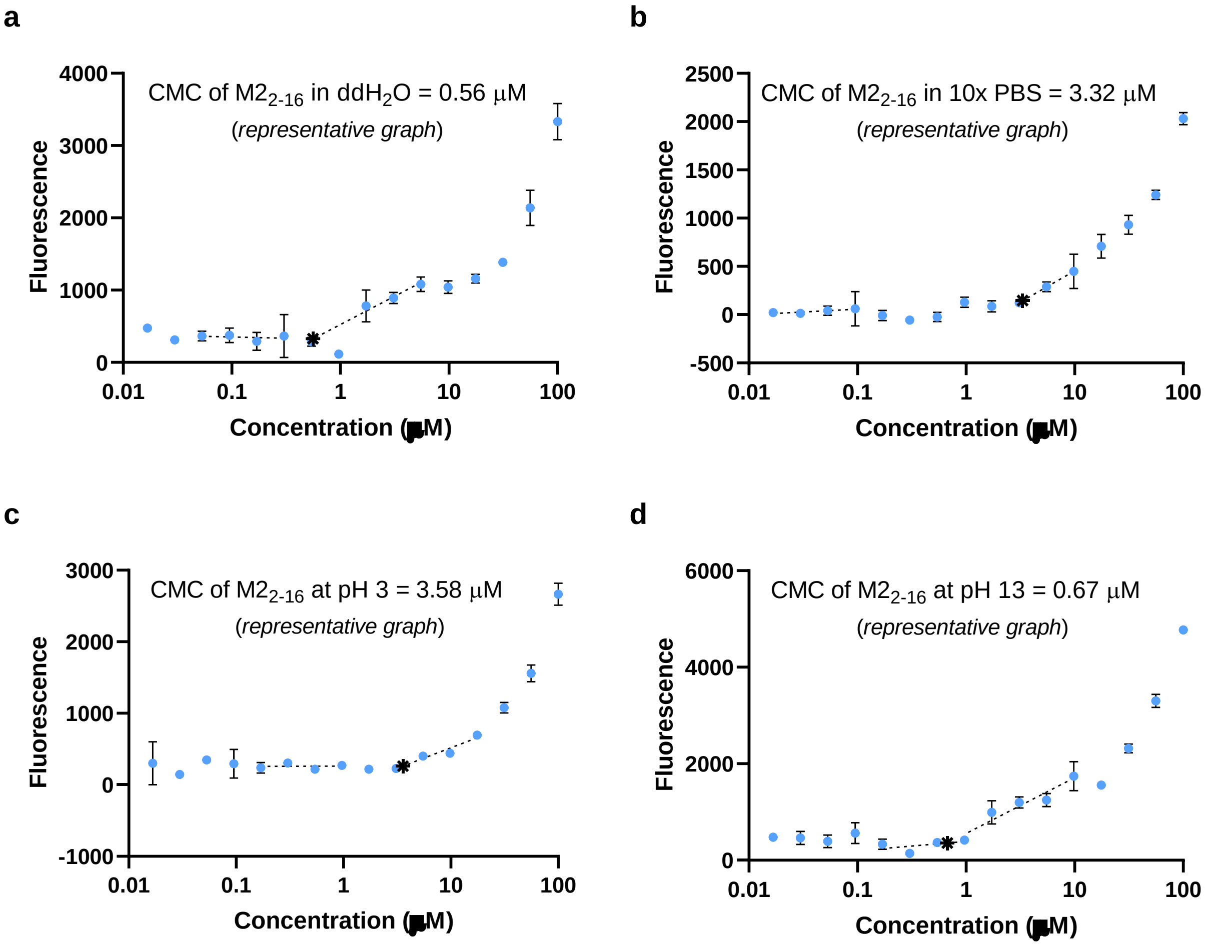


**e**

| CMC Conditions | Average CMC ± Std Dev (μM) |
|---|---|
| $ddH_2O$ | 1.47 ± 1.10 |
| 10x PBS | 2.97 ± 0.63 |
| 1x PBS at pH 3 | 3.96 ± 1.19 |
| 1x PBS at pH 13 | 2.42 ± 1.87 |

**Figure S2. Micellization of $M2_{2\text{-}16}$ peptide was not impacted by varying salt concentration nor pH.** Representative CMC graphs are shown of $M2_{2\text{-}16}$ peptide in (a) $ddH_2O$, (b) 10x PBS, (c) $ddH_2O$ at pH 3, and (d) $ddH_2O$ at pH 13. (e) CMCs of $M2_{2\text{-}16}$ peptide in the specified solution conditions are reported as the average ± standard deviation.

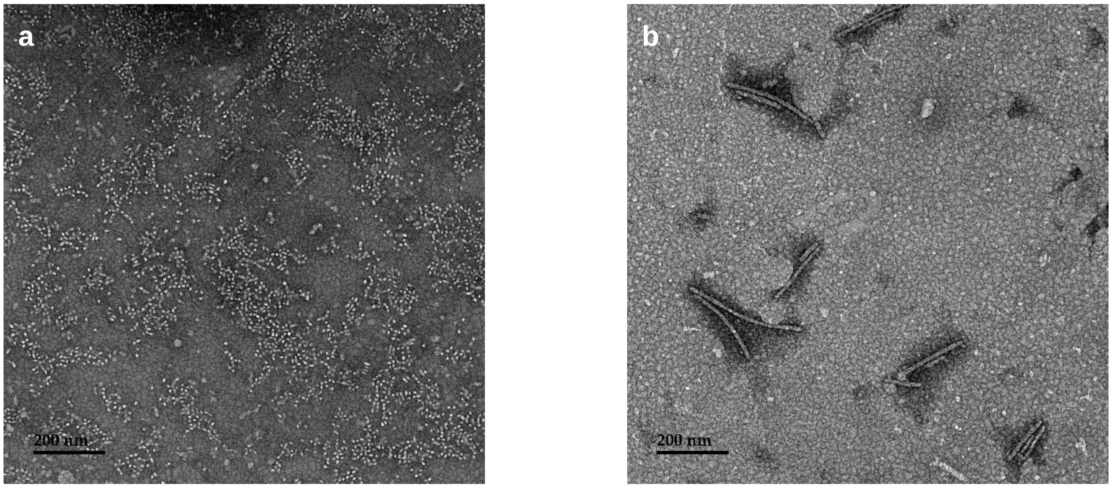


**Figure S3. $Pam_2CSK_4$ micelles had various morphologies, including (a) spheres or (b) cylinders.**

**a**

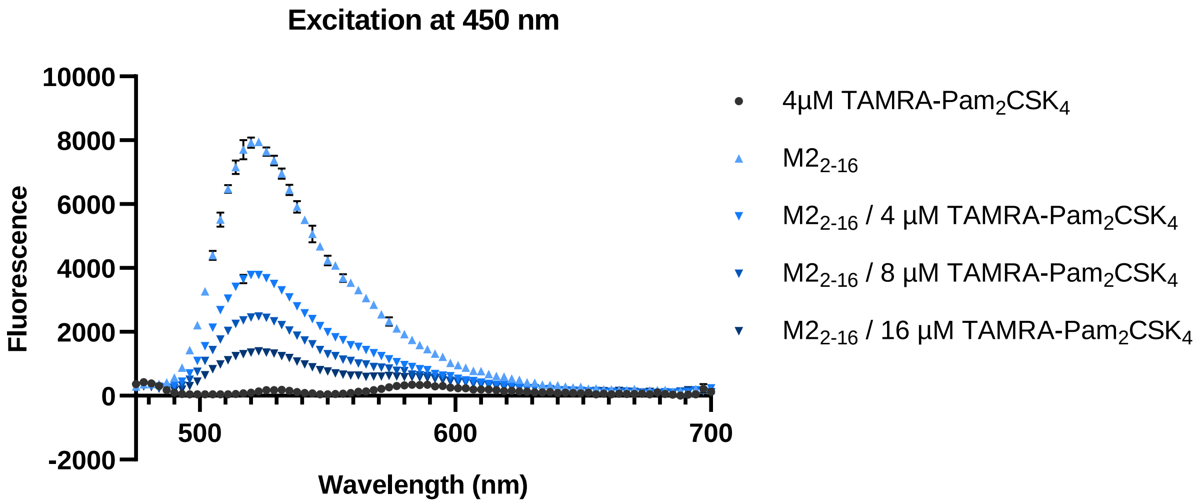


**b**

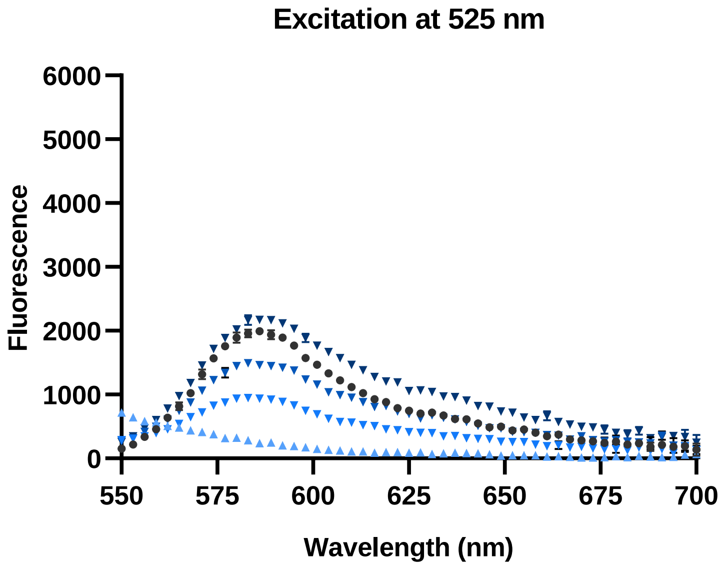


**Figure S4. Fluorescence spectra of $M2_{2\text{-}16}$ micelles with differing concentrations of TAMRA-$Pam_2CSK_4$ suggested micelle heterogeneity.** (a) Fluorescence decreased at 525 nm with increasing TAMRA-$Pam_2CSK_4$ concentration when excited at 450 nm (*i.e.*, the FAM excitation wavelength). (b) Fluorescence increased with increasing TAMRA-$Pam_2CSK_4$ when excited at 525 nm (*i.e.*, the TAMRA excitation wavelength).

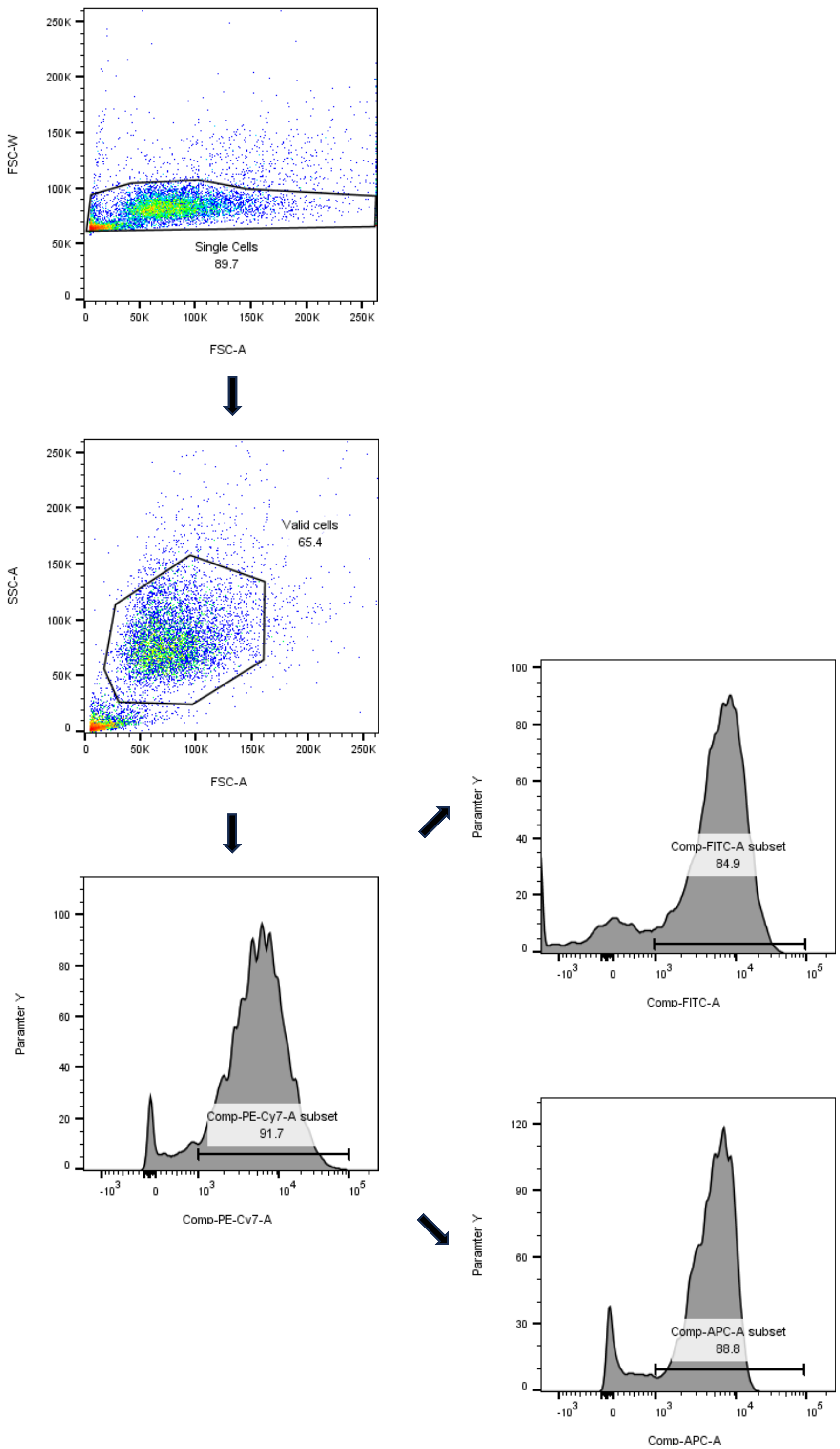


**Figure S5. Bone marrow-derived dendritic cells were gated using the strategy illustrated above.** Forward scatter (FSC-A vs FSC-W) was used to isolate out single cells (removing cell aggregates). Then cell debris was removed using FSC vs SSC (side scatter). Dendritic cells were identified by CD11c expression (PE-Cy7). Then cell activation was measured by CD40 (FITC) and MHC-II (APC) expression. Flow cytometry data was processed using FlowJo software.
